# A data-driven method to learn a jump diffusion process from aggregate biological gene expression data

**DOI:** 10.1101/2021.02.06.430082

**Authors:** Jia-Xing Gao, Zhen-Yi Wang, Michael Q. Zhang, Min-Ping Qian, Da-Quan Jiang

**Author notes:** Corresponding author, *Email address:* (Da-Quan Jiang).

## Abstract

Dynamic models of gene expression are urgently required. Different from trajectory inference and RNA velocity, our method reveals gene dynamics by learning a jump diffusion process for modeling the biological process directly. The algorithm needs aggregate gene expression data as input and outputs the parameters of the jump diffusion process. The learned jump diffusion process can predict population distributions of gene expression at any developmental stage, achieve long-time trajectories for individual cells, and offer a novel approach to computing RNA velocity. Moreover, it studies biological systems from a stochastic dynamics perspective. Gene expression data at a time point, which is a snapshot of a cellular process, is treated as an empirical marginal distribution of a stochastic process. The Wasserstein distance between the empirical distribution and predicted distribution by the jump diffusion process is minimized to learn the dynamics. For the learned jump diffusion equation, its trajectories correspond to the development process of cells and stochasticity determines the heterogeneity of cells. Its instantaneous rate of state change can be taken as “RNA velocity”, and the changes in scales and orientations of clusters can be noticed too. We demonstrate that our method can recover the underlying nonlinear dynamics better compared to parametric models and diffusion processes driven by Brownian motion for both synthetic and real world datasets. Our method is also robust to perturbations of data because it only involves population expectations.

## 1. Introduction

Previously, bulk RNA-seq technologies provide insights into complex biological systems at the population level, but they mask the heterogeneity of single cells in one population. The rapid development of single cell omics technologies makes it possible for biologists to study gene expression at the single-cell level in recent years. High-throughput and bioinformatics tools help discover cell heterogeneity, subpopulations, regulatory relationships between genes, and trajectories of distinct cell lineages. The observed heterogeneity in biological systems is driven by continuous dynamic processes (Tanay and Regev, 2017), while it remains a challenge to recover the underlying continuous time process for modeling gene dynamics directly.

Most dynamic models focus on reconstructing pseudotime trajectories. Saelens et al. (2019) compared 45 kinds of trajectory inference (TI) methods on 110 real and 229 synthetic datasets, but there is no consensus that what is the best tool (Luecken and Theis, 2019). These algorithms that utilize the continuity of single cell data to reconstruct trajectories may induce model bias. Meanwhile, RNA velocity based methods (La Manno et al., 2018; Svensson and Pachter, 2018; Bergen et al., 2020; Li et al., 2020) are burgeoning in recent years. However, the assumptions in their models are not applicable in all systems. For instance, hematopoietic stem cells (HSCs) retain more of the unspliced mRNAs and get more spliced mRNAs under activation (Bowman et al., 2006). Another approach based on optimal transport analysis can also be used to recover trajectories (Schiebinger et al., 2019).

Stochastic differential equations (SDEs) are feasible solutions to model the stochastic dynamics of gene expression data. For biochemical systems, a series of ordinary differential equations are often used to describe the evolution of reactant concentrations following the law of mass action (Keener and Sneyd, 2009). However, for the lack of randomness, transitions among multiple attractors are infeasible to take place for multistable systems. The inherent stochasticity observed in many important cellular processes, such as transcription and translation, attracts scientists to interpret biological phenomena using stochastic models. Niu et al. (2016) used reflected SDEs to model the biochemical reaction systems. Jia et al. (2014) explained the underlying molecular mechanisms of phenotype switching in bacteria by a nonlinear SDE. Forman and Sørensen (2014) estimated the folding and unfolding rates of the small Trp-zipper protein using multi-modal diffusions. Except for SDEs driven by Brownian motion with continuous trajectories, more general Lévy-driven SDEs are introduced (Applebaum, 2009). Jia et al. (2017) deduced that the macroscopic limit of the protein number is a switching Lévy-driven SDE, and interpreted the existence of protein burst phenomena due to the imbalanced decay rates of mRNA and protein.

In this contribution, we propose using a jump diffusion process rather than a continuous diffusion process driven by Brownian motion to model the dynamic evolution of gene expression in the single-cell resolution. Compared to diffusion processes which are solutions to SDEs driven by Brownian motion with Markovianity, our jump diffusion process contains both Brownian motion and a compound Poisson process. There are three reasons for our choice. First, large jumps caused by the compound Poisson process benefit transitions among attractors. In a multistable system, the transition from one attractor to another non-adjacent one can be accomplished by a compound Poisson process, while Brownian motion can not induce this phenomenon. Second, jump diffusion processes can fit biological systems with non-Gaussian distributions. For example, the bursty and intermittent production of mRNA and proteins creates variation in individual cells and causes heavy tailed distributions (Zheng et al., 2016). Third, when the jump intensity is small or jump amplitude near to zero, our model degenerates to a diffusion process. Thus, our model is a generalization of diffusion processes driven by Brownian motion. Nowadays, SDEs with Lévy noise are widely used in modeling gene expression (Xu et al., 2016; Jia et al., 2019; Cai and othres, 2019; Chen and Jia, 2020) and the interpretation of burst behavior from a mathematical perspective can be referred in Bokes et al. (2012) and Jia (2017).

The reconstruction of complex nonlinear dynamic systems using SDEs driven by Brownian motion is not new. For parametric models, the drift and diffusion coefficients are assumed to have particular forms. For example, the Black-Scholes model with linear coefficients is applied to price an option in finance. SDEs with Gaussian mixture potentials are used to model the movement of animals in ecology (Preisler et al., 2013; Gloaguen et al., 2018). For such models, one only needs to determine finitely many parameters based on experimental data, but suffers from their limited model capacity. Correspondingly, high expressive models parameterize the drift coefficient to be a neural network and diffusion coefficient to be a constant (Hashimoto et al., 2016; Wang et al., 2018; Ma et al., 2020). On the other hand, Tabar (2019) discussed data-based reconstruction procedures, but their estimator involves a conditional averaging of a small time limit over Brownian trajectories, which can only be applied to high-resolution time series data. While in practice, measuring the gene activity of individual cells involves destroying the cells so that their content can be analyzed. Therefore, we are unable to acquire the information of each cell all the time due to technical limitations. We refer to observations made in these scenarios as aggregate data.

Our model aims to learn a jump diffusion process from aggregate gene expression data based on real time rather than pseudotime. The state variables are the expression levels of genes in a single cell, and the coefficients for the jump diffusion process are modeled as neural networks. Once the jump diffusion process is achieved, its trajectories can be taken as the evolution of cells. The stochasticity brought by Brownian motion and compound Poisson processes determines the heterogeneity of cells. Given the current state of a cell, its subsequent states are also stochastic. The average instantaneous rate of state change reveals the next step evolution, thus can be taken as “RNA velocity”. As can be seen from our experimental results, our method can model the cell dynamics more accurately and is robust to perturbations of data. Also, from the proposed method we derive and interpret three vital biological concepts, i.e. cluster, velocity, and trajectory.

## 2. Material and methods

### 2.1. Dataset

Assuming that we have observed totally *D* time points during the whole time interval [0, *T*], where 0 = *T*_0_, *T*_1_, ···, *T*_*D*–1_ = *T* and the time partition may not be equal. At each of these time points, there are *N_i_* independent and identically distributed (*i.i.d*.) samples 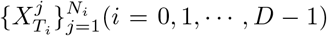 that we term aggregate observations. The individuals 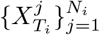 observed at time *T_i_* are often not identical to those 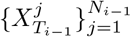 observed at the previous time *T*_*i*–1_. Since the full trajectory of each individual is not available, the only useful information at time *T_i_* is its distribution, written as *P*(*x, i*). While, in practice, we can just achieve the empirical distribution 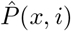 at time *T_i_* from 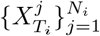 as an alternative.

For example, Klein et al. (2015) studied the differentiation of mouse embryonic stem cells. The expression levels of 24,175 genes for 933, 303, 683, and 798 cells at Day 0, Day 2, Day 4, and Day 7 are quantified respectively. In this case, we have *D* = 4, *T*_0_ = 0, *T*_1_ = 2, *T*_2_ = 4, *T*_3_ = 7 and *N*_1_ = 933, *N*_1_ = 303, *N*_2_ = 683, *N*_3_ = 798.

### 2.2. Wasserstein distance

Aggregate gene expression data, which are snapshots of a continuous time cellular process, can be regarded as *D* empirical distributions. The discrepancy between empirical distributions and predicted distributions by our jump diffusion process is measured by Wasserstein distance for its smoothness (Arjovsky et al., 2017). Wasserstein distance is also used in parameter estimation of biochemical reaction networks (Öcal et al., 2019). Wasserstein-1 distance, also called Earth-Mover distance, is defined by

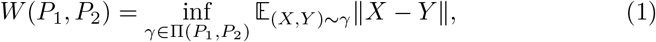

where Π(*P*_1_, *P*_2_) denotes the set of every joint distribution *γ* whose marginals are *P*_1_, *P*_2_ respectively. In practice, Formula (1) is hard to use, so Villani (2008) gave its duality

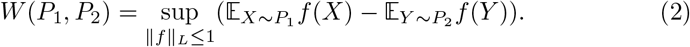

Here, ║*f*║_*L*_ ≤ 1 denotes that *f* is 1-Lipschitz continuous.

### 2.3. Algorithm

#### 2.3.1. The jump diffusion model

Aggregate single-cell gene expression data are modeled as outputs of a discontinuous jump diffusion process. Namely, the evolution of individual cells satisfies

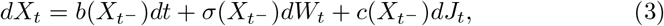

for *t* ∈ [0, *T*]. The process is defined on a filtered probability space 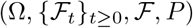. *X_t_* is a *d*-dimensional state variable representing the expression levels of *d* genes in a cell. 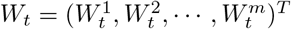 denotes a standard *m*-dimensional Wiener process. 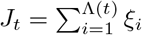 is a compound Poisson process where Λ = {Λ(*t*): *t* ≥ 0} is a Poisson process with intensity λ, {*ξ_n_*: *n* ≥ 1} are *i.i.d*. random variables independent of the Poisson process Λ, and *ξ_n_* denotes the *n*th jump amplitude with density function 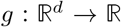. The two processes *W_t_* and *J_t_* are mutually independent. We also assume 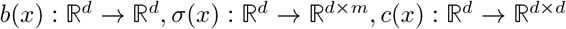. Coefficients *b*(*x*), *σ*(*x*) and *c*(*x*) are set to be state dependent only. The undetermined parameters are denoted by 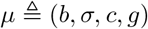.

#### 2.3.2. Predicted distributions by the jump diffusion process

Here, we take the first two time points *T*_0_ and *T*_1_ as an example to show how we predict the distribution at *T*_1_ given the empirical distribution at *T*_0_. The time interval between *T*_0_ and *T*_1_ is far from being sufficiently small, so we construct an equidistant time discretization with *T*_0_ = *t*_0_, *t*_1_, ···, *t*_*n*_1__ = *T*_1_. The most widely used Euler-Maruyama (EM) discretization scheme that solves Equation (3) numerically is given by

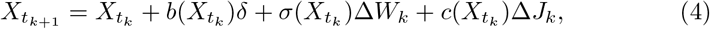

where *δ* = (*T*_1_ – *T*_0_)/*n*_1_, Δ*W_k_* = *W*_*t*_*k*+1__ – *W_t_k__* and Δ*J_k_* = *J*_*t*_*k*+1__ – *J_t_k__*. That is to say, *N*_0_ samples evolve *n*_1_ steps from *T*_0_ to *T*_1_ following Equation (4), and the generated samples at *t_j_*(*j* = 1, 2, ···, *n*_1_) are denoted as 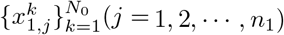. In this way, we achieve predicted distribution 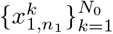 at *T*_1_, written as 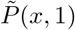. Similarly, we can achieve 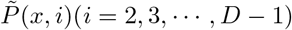

#### 2.3.3. Cost function

The dynamics can be learned by minimizing the Wasserstein distance between the empirical distribution 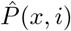 and the predicted distribution 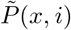. Our cost function can be chosen to be 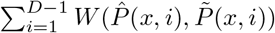. Therefore, the task is to solve the following min-max problem

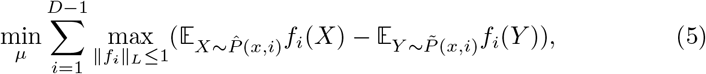

which is similar to Wasserstein generative adversative nets (WGAN). The difference is that we have totally *D* – 1 critics *f_i_*(*i* = 1, ···, *D* – 1). The overview of our algorithm is shown in Figure 1.

**Figure 1:**
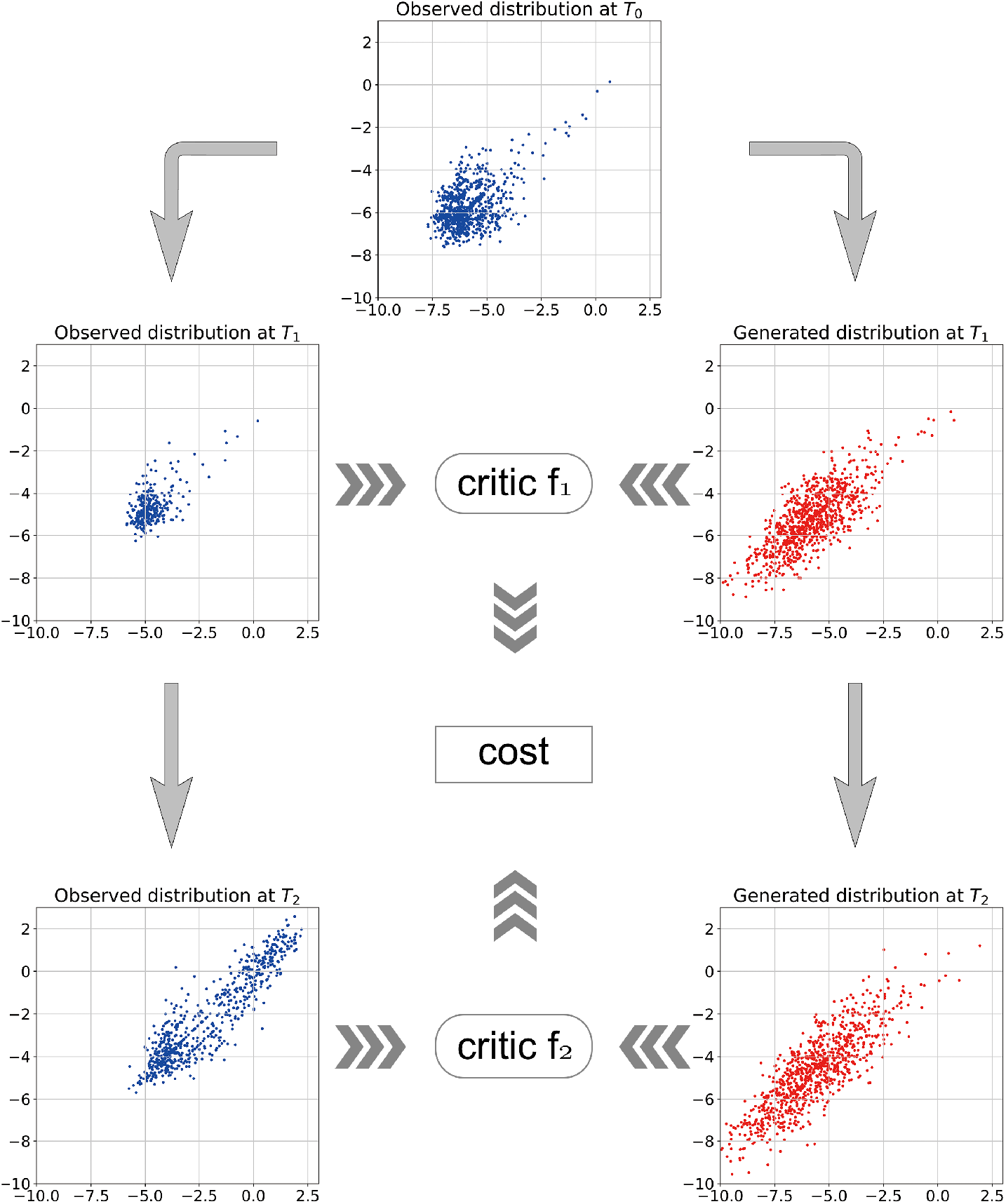
The overview of our algorithm for learning stochastic dynamics from aggregate data. The left side has three observed distributions (blue points), and the right side has two predicted distributions (red points) by Equation (4). At time *T*_1_ or *T*_2_, the critic function takes the input of an observed and a predicted distribution and outputs a scalar as the Wasserstein distance whose summations at all time points will be our cost.

In Formula (5), there are two expectations to compute. The first one is easy, and the formula is

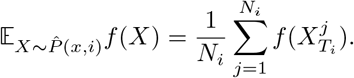

For the second one, after achieving predicted samples by Equation (4) at *T_i_*, we have

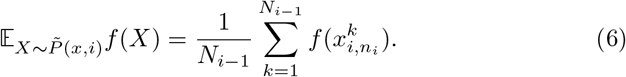

Recall that 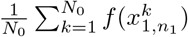 is an estimator of 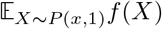 for real distribution *P*(*x*, 1) at *T*_1_ and Lipschitz continuous observables *f*. Its mean squared error (MSE) has the following result.

##### Theorem 1.

*For the test function f satisfying* ║*f*║_*L*_ ≤ 1, *the* MSE *of* 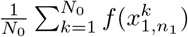 *is asymptotically O*(1/*N*_0_) + *O*(*δ*).

#### 2.3.4. Training framework

In a cellular process, genes that are not informative will be filtered out. For such a gene, the distribution at final time point *T*_*D*–1_ will have small difference with the one at the starting time point To. Therefore, we sort genes by the Wassserstein distance between *T*_*D*–1_ and *T*_0_, and genes with higher Wasserstein distance will be selected (Hashimoto et al., 2016).

*b*(*x*), *σ*(*x*), *c*(*x*) and *f_i_*(*i* = 1, 2, ···, *D* – 1) are unknown functions, so multilayer perceptrons are used to model these functions. *g*(*x*), as a probability density function, can be chosen to be Gaussian, see section 3.1. WGAN framework is used to solve the min-max problem 5. The evolution formula (4) takes the role of generator *G* with parameter *μ*. WGAN-div (Wu et al., 2018) is used to implement the 1-Lipschitz constraint of critic functions *f_i_*. Let *P_u_* be the mixed distribution of 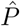 and 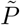, and *k, p* are hyperparameters used in Wu et al. (2018) with recommended values *k* = 2, *p* = 6. We give the pseudo code of our algorithm in the following.

**Algorithm 1.**
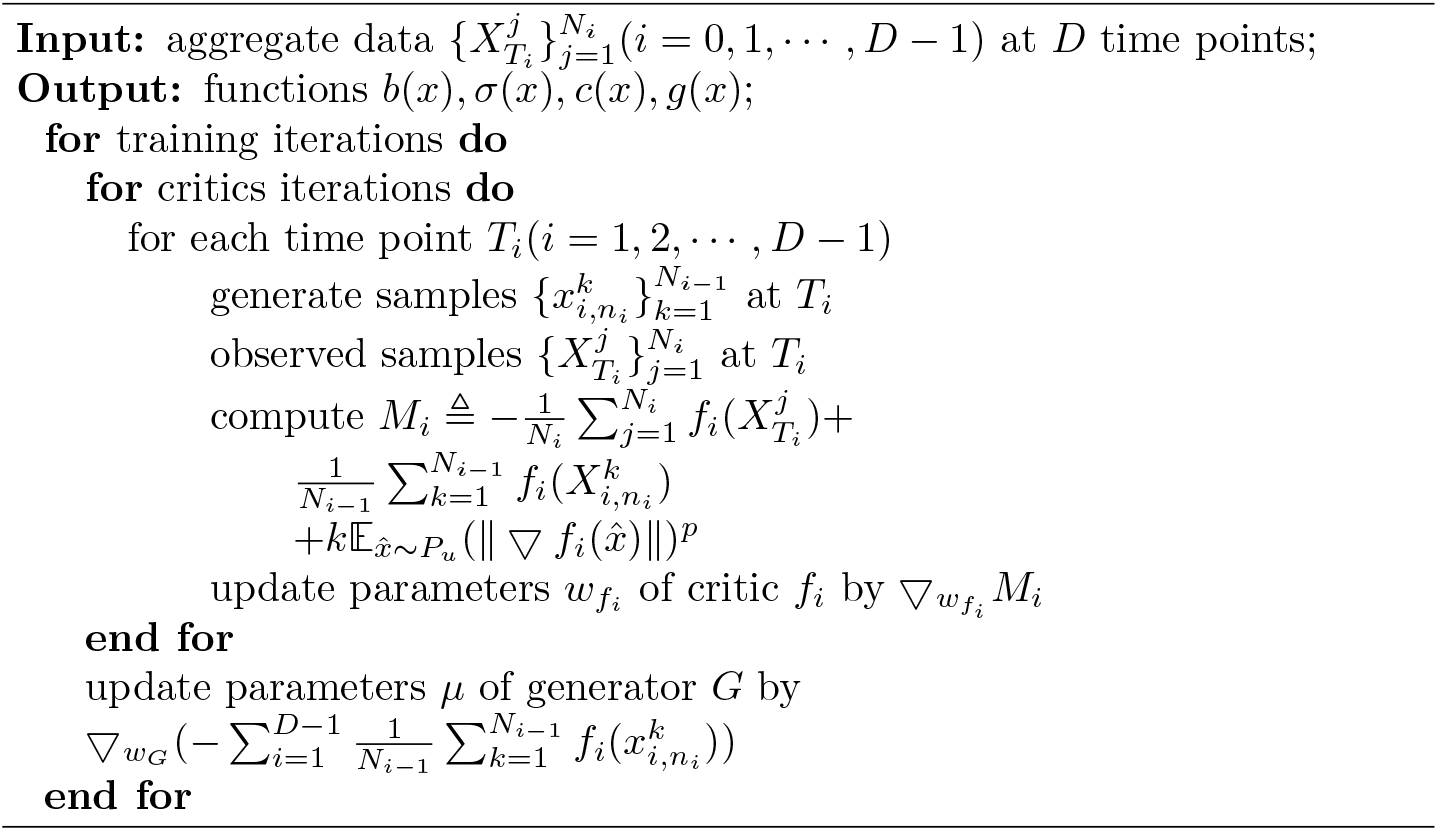
Reconstruction of nonlinear dynamics (3) from aggregate observations.

## 3. Results

We compare the performance of our jump diffusion process (3) with OU process and diffusion processes driven by Brownian motion. The OU process is the solution to *dX_t_* = *θ*(*μ* – *X_t_*)*dt* + *σdW_t_*, and is a stationary Gauss–Markov process. As an example of parametric models, it has a wide range of applications in financial mathematics and physics. Diffusion processes driven by Brownian motion *dX_t_* = *b*(*X_t_*)*dt* + *σdW_t_* can model nonlinear dynamics and have a higher model capacity than OU process.

### 3.1. A synthetic dataset

We first evaluate our algorithm on a synthetic dataset generated by the following diffusion process

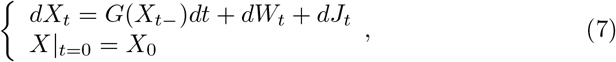

where *G*(*x*) has the following form

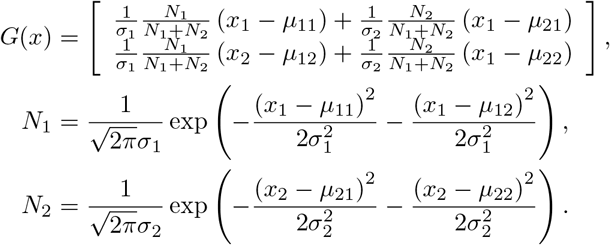

Parameters are set to be *μ*_1_ = (16,12), *μ*_2_ = (–18, –10), (*σ*_1_, *σ*_2_) = (1, 0.95), λ = 40 and jump size *ξ* = (*ξ*_1_, *ξ*_2_), where *ξ*_1_, *ξ*_2_ are independent exponential random variables with means 0.1 and 1, respectively. The initial distribution for *X*_0_ is a standard Gaussian with mean (0,0) and covariance **I**_2_. Starting from *X*_0_, Equation (7) evolves following EM scheme (4) with *δ* = 0.02. We utilize 1,200 simulated samples at time 0*δ*, 3*δ* and 6*δ* as training sets and predict distributions at 2*δ*, 4*δ* and 10*δ*.

We compare the performance of the OU process, diffusion processes driven by Brownian motion (denoted as SDEnoJump), and our jump diffusion model. Our model with exponential jump size is written as SDEexpJump and Gaussian jump as SDEgauJump. As can be seen in Figure 2, the training process for OU does not converge within 40,000 epochs, and the Wasserstein loss at time 3δ keeps increasing. SDEnoJump works better. Its Wasserstein loss is lower than OU but still can not compete with our jump models. For the underlying jump size follows a 2-dimensional exponential distribution, SDEexpJump converges much faster and more stably than the other three methods. Although SDEgauJump converges slower than SDEexpJump, it can achieve good results too.

**Figure 2:**
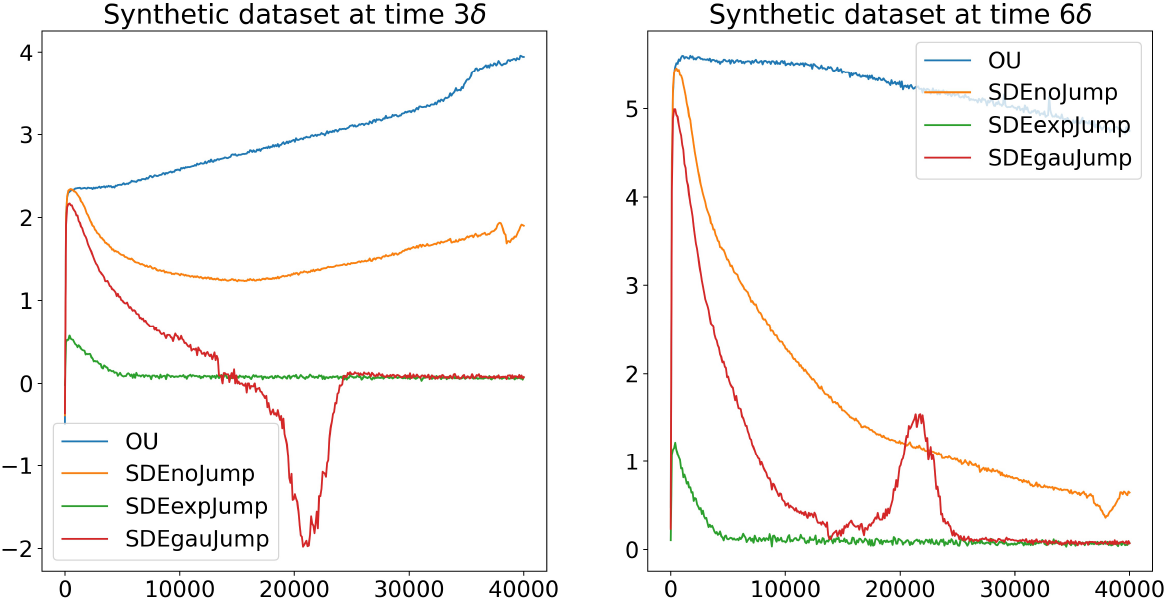
Wasserstein loss v.s. iterations for the synthetic dataset at two observed time points *3δ* and 6δ. SDEexpJump and SDEgauJump perform better because their Wasserstein distance is close to 0.

The predicted distributions by the four methods at 2*δ*, 4*δ*, 10*δ* are shown in Figure 3. The Sinkhorn distances, which are approximations of Wasserstein distance and easy to compute, are displayed in Table 1. OU performs the worst due to its small capacity. SDEnoJump can learn the dynamics, but Figures 3(e) and 3(f) show that the upper right and bottom left regions are worse predicted. SDEexpJump and SDEgauJump both can generate good results, so we will use SDEgauJump in the following experiments for the need of vectorization. Also, from the experiments, we conclude that diffusion processes driven by Brownian motion can not recover the dynamics from the data generated by a jump diffusion process while jump diffusion processes can recover the dynamics from the data generated by diffusion processes driven by Brownian motion.

**Figure 3:**
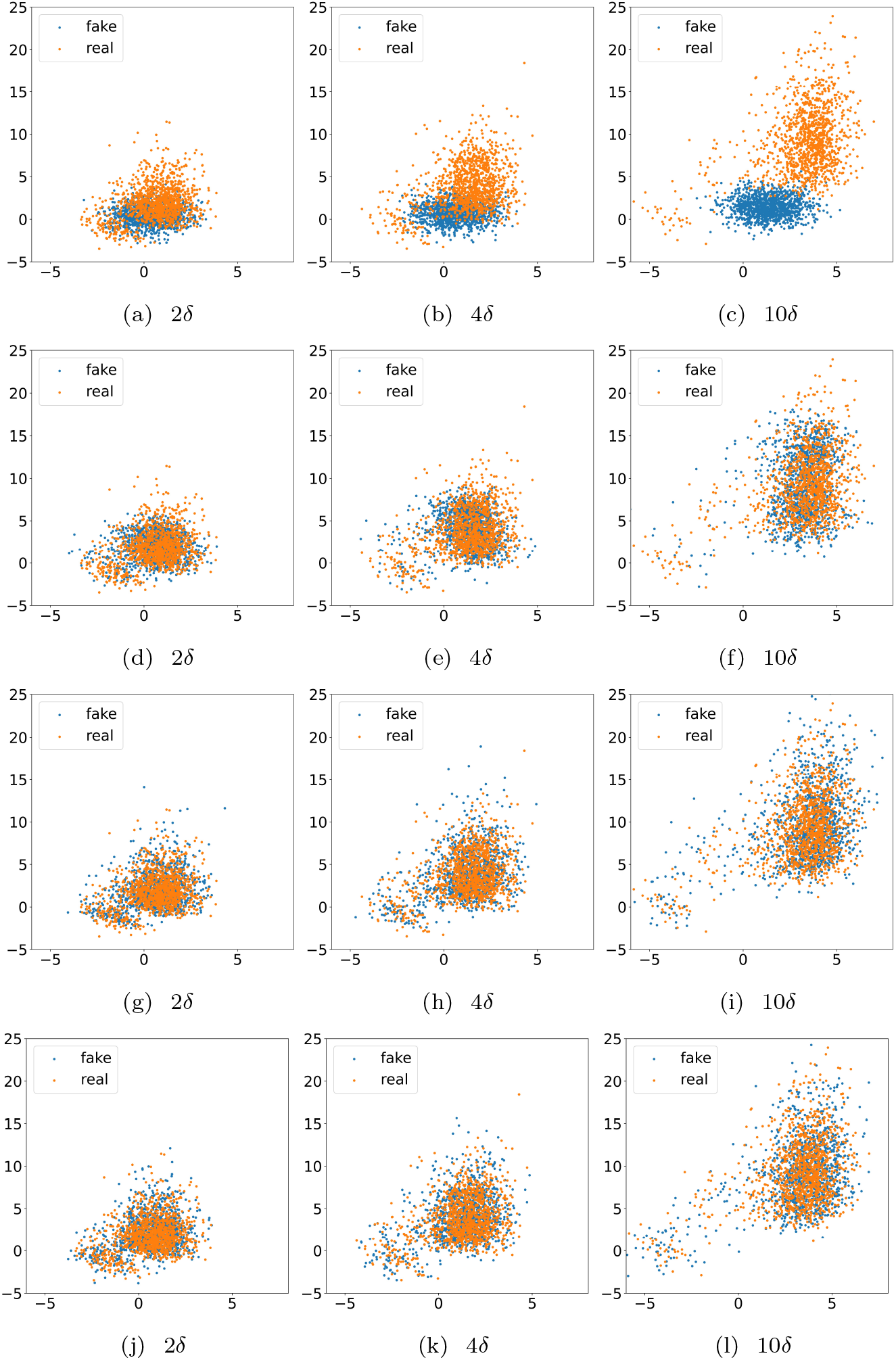
The real are observed distributions and the fake are predicted distributions in a 2-dimensional space. The performance of four models, i.e. OU((a) to (c)), SDEnoJump((d) to (f)), SDEexpJump((g) to (i)) and SDEgauJump((j) to (l)) at three time points 2*δ*, 4*δ*, 10*δ* is compared.

**Table 1:** The Sinkhorn distances between true distributions and predicted distributions of different models on the synthetic dataset.

| Model | OU | SDEnoJump | SDEexpJump | SDEgauJump |
| --- | --- | --- | --- | --- |
| $2\delta$ | 1.7411 | 0.1717 | 0.0122 | 0.0325 |
| $4\delta$ | 6.4121 | 0.1777 | 0.0429 | 0.0002 |
| $10\delta$ | 36.4112 | 0.2168 | 0.0445 | 0.0873 |

### 3.2. Stem cell differentiation dataset

In this part, we use the jump diffusion process to model the development of embryonic stem cells (Klein et al., 2015). These cells differentiate from embryonic stem cells to functional cells after the removal of LIF (leukemia inhibitory factor) at day 0 (D0). The expression levels of 24,175 genes for 933, 303, 683, and 798 cells at D0, D2, D4, and D7 are quantified respectively. We select the top ten variables determined by the Wasserstein distance between D0 and D7 distributions, and they are *Krt8, Krt18, Tdh, Tagln, Mt1, Dppa5a, Mt2, Gsn, Lgals1, and Pou5f1*. The preprocessing procedures, i.e. normalization and imputing missing expression levels, are the same as that in Hashimoto et al. (2016).

#### 3.2.1. Prediction

We evaluate our algorithm using two prediction tasks corresponding to middle states and terminal states of a dynamic process. The first task is to predict the cell distribution at D4 given aggregate observations at D0, D2, and D7, and the second is to predict D7 based on D0, D2, and D4. Wang et al. (2018) compared the performance of their LEGEND with OU and NN (Hashimoto et al., 2016) in predicting *Krt8* and *Krt18* at D4. As reported in their paper, none of the above three methods can predict the bimodal distribution of *Krt8* in the ground truth at D4. Therefore, it is necessary to consider a more complex jump diffusion process with high capacity to model these biological dynamics.

We first show in Table 2 that compared to OU and SDEnoJump (Hashimoto et al., 2016; Wang et al., 2018; Ma et al., 2020) our jump diffusion model achieves the lowest Sinkhorn distances in both two predicting tasks. In predicting *Krt8* at D4, our model is the only one that successfully learns the bimodal behavior. Figures 4 and 5 also show that our predicted distributions are much closer to the underlying true distributions, implying that our model can actually recover the dynamic behavior from aggregate gene expression data.

**Table 2:** The Sinkhorn distances between empirical distributions and predicted distributions of different models on the stem cell differentiation dataset.

| Model | OU | SDEnoJump | SDEgauJump |
| --- | --- | --- | --- |
| $D4$ | 10.0763 | 5.8835 | 3.5898 |
| $D7$ | 15.7564 | 16.2900 | 7.3074 |

**Figure 4:**
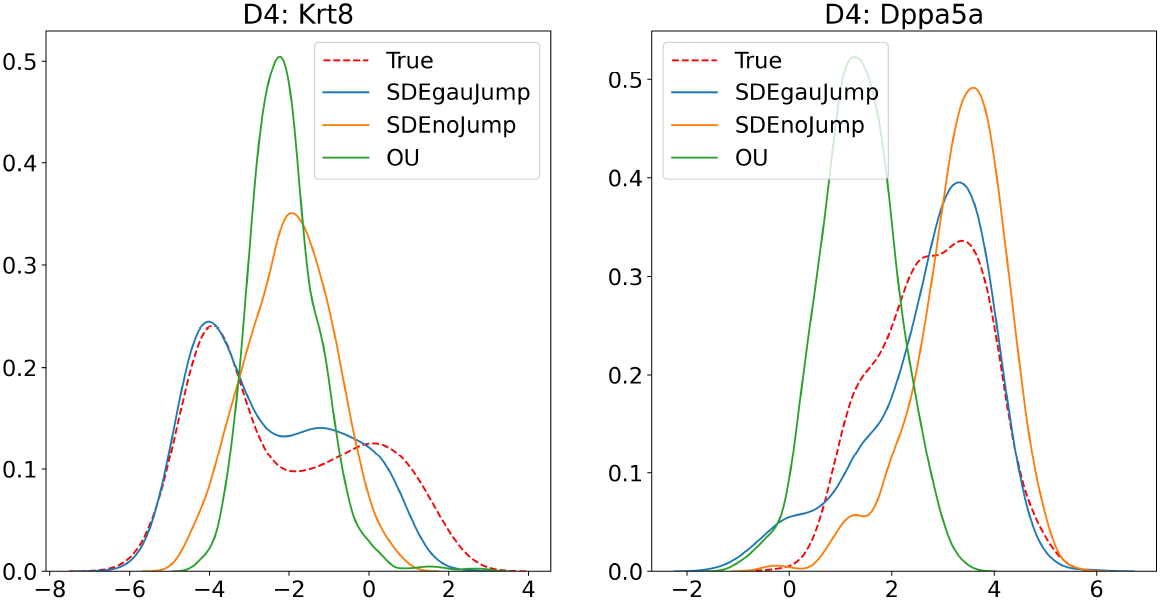
Lines are probability density functions estimated by kernel density estimation. The red dotted lines are achieved by the observed data of Krt8 and Dppa5a at D4 and the solid lines by the predicted samples of Krt8 and Dppa5a at D4.

**Figure 5:**
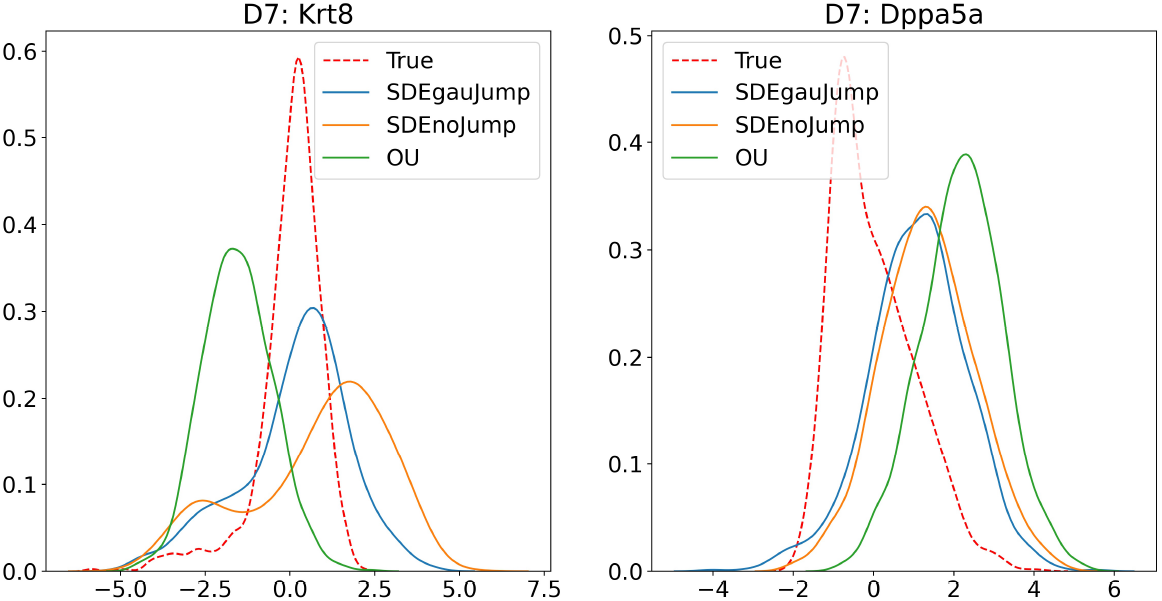
The red dotted lines are achieved by the observed data of Krt8 and Dppa5a at D7 and the solid lines by the predicted samples of Krt8 and Dppa5a at D7.

#### 3.2.2. Evolution

Once the jump diffusion process is achieved, the learned model can be utilized to calculate or predict some meaningful quantities in a biological process. Therefore, we use complete gene expression data at D0, D2, D4, and D7 to learn the underlying dynamics. The predicted distributions by our learned model match the observed empirical distributions perfectly at D2, D4, and D7 (Supplementary Figures 1, 2, and 3). PCA (Principal Component Analysis) and tSNE with two components are applied to visualize the learning results. The predicted and true distributions match quite well, and the indistinguishable images enable us to believe that gene expression data actually evolve following our estimated jump diffusion dynamics no matter which dimension reduction method is used. Points on the right side of Figure 6 are denser than those on the left because we generate 933 trajectories from D0. In contrast, we should be careful about results reported by Monocle (Figure 7) and other pseudotime trajectory inference methods. We model gene dynamics using real observed time not pseudotime.

**Figure 6:**
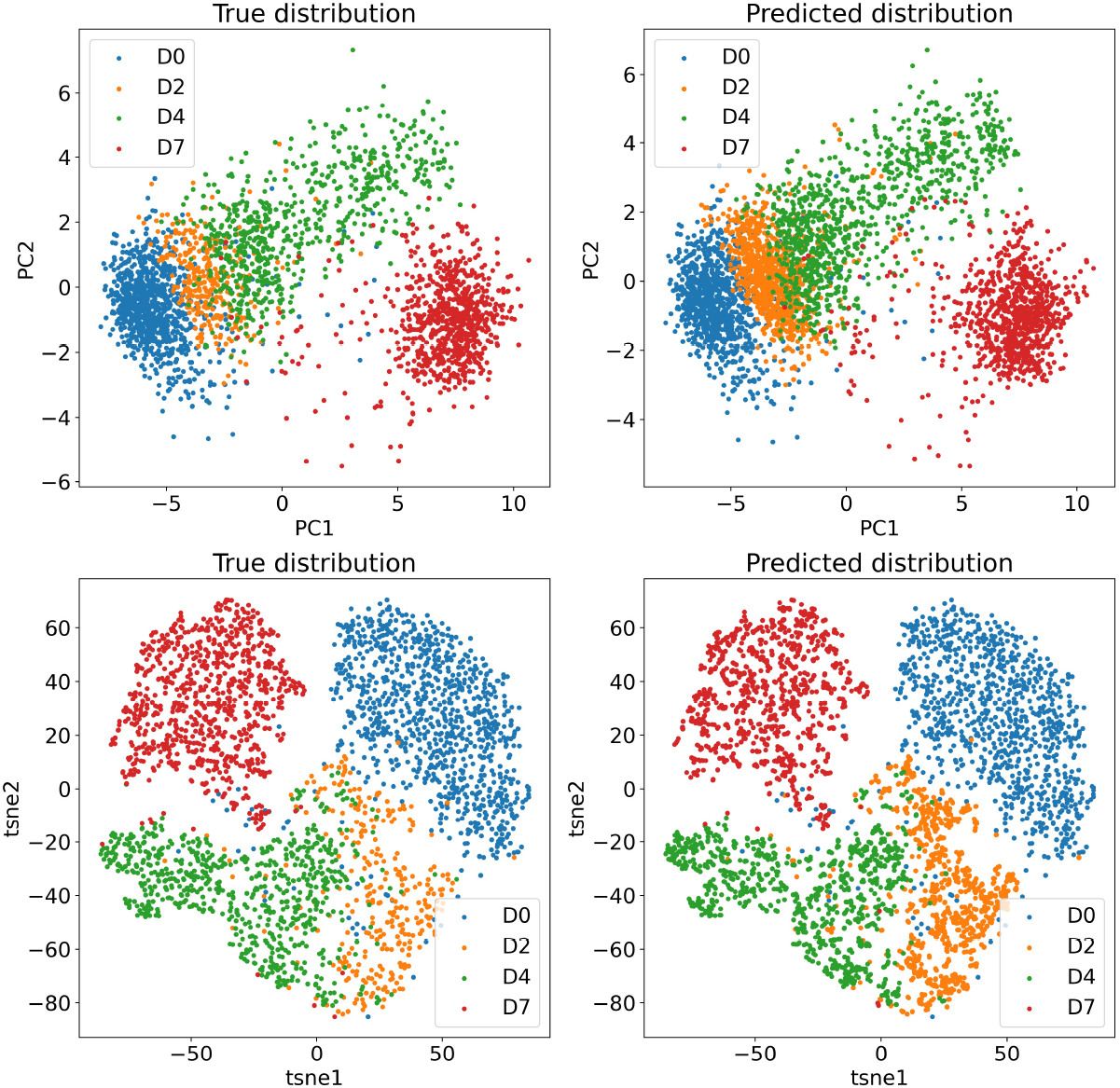
PCA and tSNE are applied to visualize our learning results. The left side shows the true distributions at four time points and the right shows the corresponding predicted distributions.

**Figure 7:**
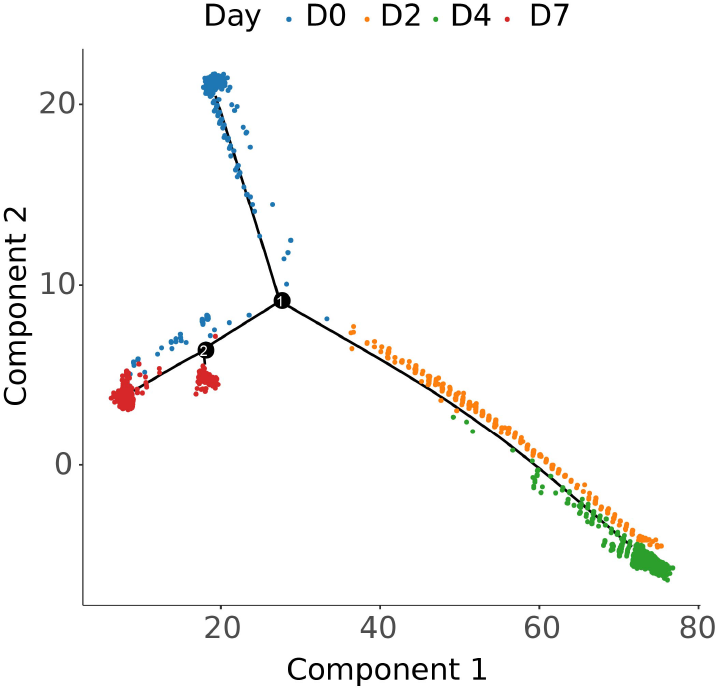
Trajectory inferred by Monocle with DDRTree method.

#### 3.2.3. Subpopulation trajectory heterogeneity

For Equation (3), its solution will be a stochastic process *X_t_*(*t* ≥ 0). If we fix *ω* ∈ Ω, *X_t_*(*ω*) is a function of *t* which corresponds to the gene expression changing with t. Therefore, the development of a cell can be seen as a realization of the jump diffusion process. For there are 933 cells at D0, we have totally 933 trajectories with different trends. However, to get more robust results, we focus on subpopulations rather than a single trajectory.

To understand the biological process better, we analyze the lineage-specific gene dynamics during the development process. *Dppa5a*, an important gene involved in the maintenance of embryonic stem cell pluripotency, shows different dynamics in trophectoderm, primitive endoderm, and epiblast (Petropoulos et al., 2016). In Figure 8, three subpopulation trajectories with different behavior are picked out by K-means for time series data. The predicted trend of *Dppa5a* is coincident with the results in Petropoulos et al. (2016). The average expression level of *Krt8* also agrees with the outcome in the original paper (Klein et al., 2015). These suggest our method could model the lineage-bias of individual cells.

**Figure 8:**
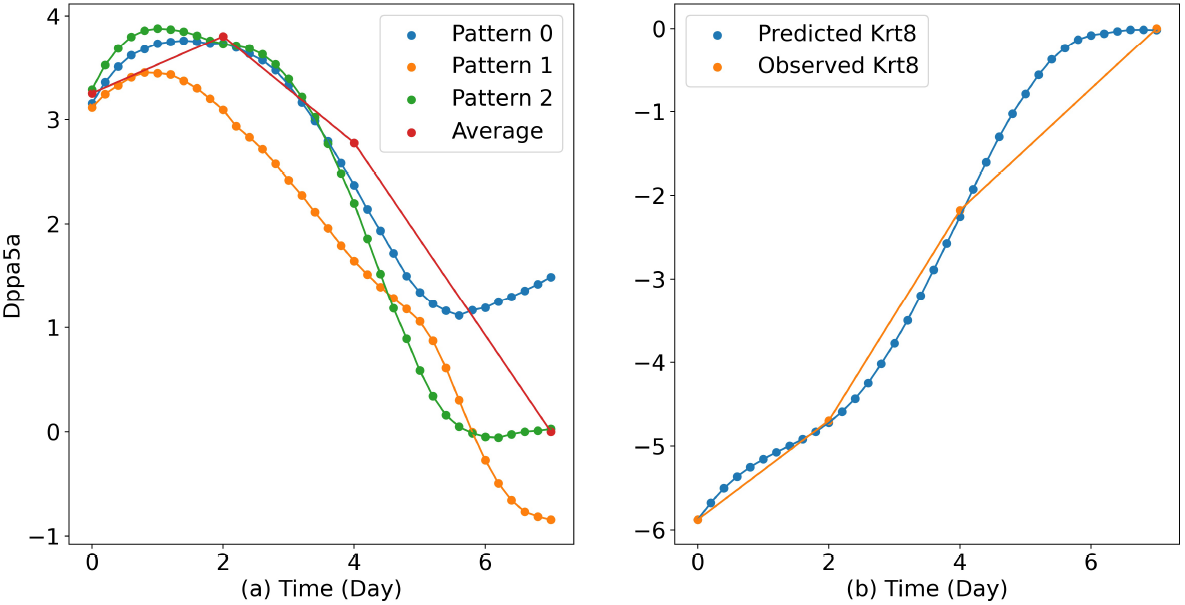
(a) Three clusters with different trend patterns of the expression levels of Dppa5a are picked out. The red points represent the average expression level at four observed time points. (b) Predicted and observed average expression levels of Krt8.

#### 3.2.4. RNA velocity

The next step evolution of cells differs not only in direction, but also in speed. RNA velocity is just a quantity that measures the instantaneous speed and direction of motion. Based on the current state, RNA velocity describes the direction and speed of state transition. It is uncorrelated to the previous time, so the Markovian property of our jump diffusion (3) can be utilized to compute RNA velocity. Given the current state *X_t_k__* at time *t_k_*, the expected state at *t*_*k*+1_ is given by

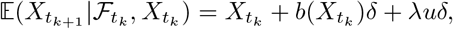

where *u* is the mean of jump size *ξ*. Thus, the expected velocity at *t_k_* is given by

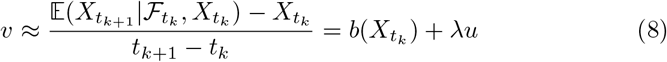

In Table 3, we compute the average size of RNA velocity at four time points. Cells at D4 evolve much faster than those at D2 and D7 because lineage-specific genes increase in variability and proliferation rate (Petropoulos et al., 2016; Waisman et al., 2019). The next step evolution of individual cells in Figure 9 offers the evidence for short-time cell developmental trajectories. These results demonstrate that our method could model the long-time and short-time dynamics of single cells simultaneously.

**Figure 9:**
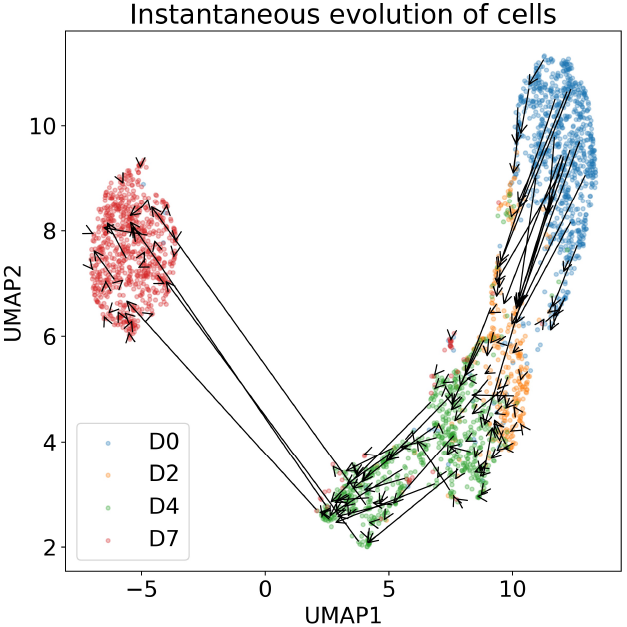
The next step evolution for some cells visualized by the UMAP dimension reduction method with two components. The points at four days are real data.

**Table 3:** Average size of *v* at four days.

| Time | D0 | D2 | D4 | D7 |
| --- | --- | --- | --- | --- |
| $v$ | 21.7149 | 12.1285 | 26.7343 | 14.6009 |

#### 3.2.5. Clusters

Clusters change in scales and orientations in the cell differentiation process. Without losing generality, we select *Krt8* as an example to show that. It can be observed from the experimental data that one cluster moves flatly from D0 to D2. Two clusters appear at D4 and merge to one at D7 finally. We use our jump diffusion model to supplement the intermediate development information (Supplementary Figure 4). The situations at two time points D4.5 and D5, which could be transition states, are predicted by our method (Figure 10). Two clusters have nearly the same size at D4.5, and the right cluster is in the leading position at D5.

**Figure 10:**
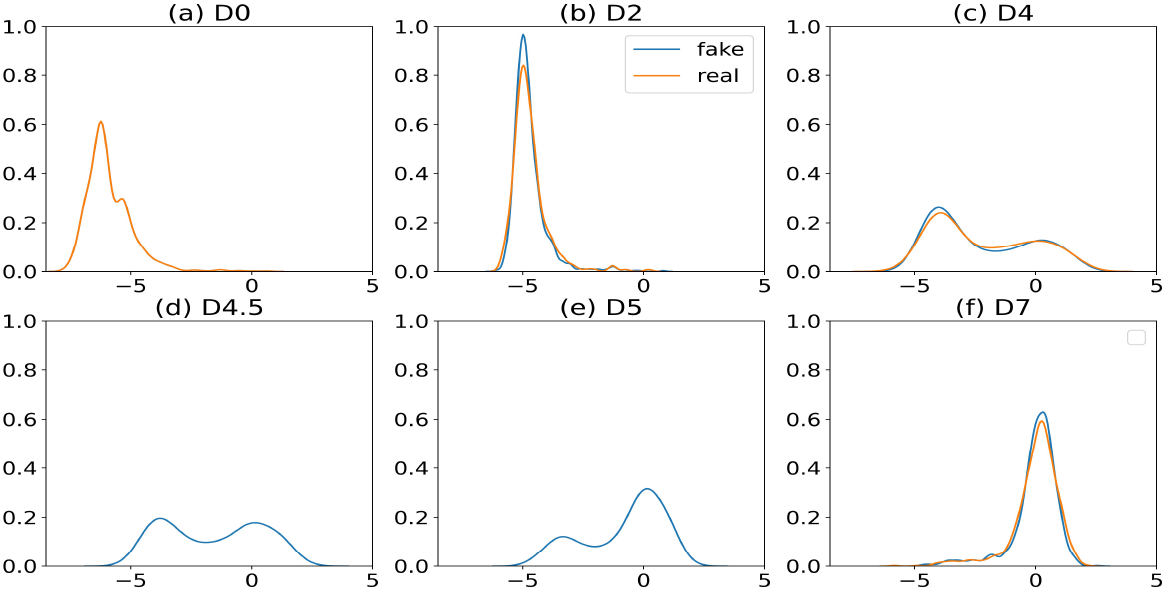
Lines are plotted by kernel density estimation. The Krt8 expression changes with time from (a) to (f). In (a), there is only one observed distribution because D0 is the starting time point. (d) and (e) are predicted by our method.

### 3.3. Cell cycle dataset

We show our method can reveal non-equilibrium biological processes. 280 mouse ESCs in Buettner et al. (2015) are studied, and three stages G1, S, and G2M of the cell cycle have 96, 88, 96 cells respectively. PHATE (Moon et al., 2019) is used to reduce the data into a 2-dimensional space. We use the OU process, SDEnoJump and SDEgauJump to model the cell cycle dynamics. The OU process drives cells to go to the center as expected because the central region can be taken approximately as an average of the three stages so as to achieve lower Wasserstein distance, see Figure 11. Figure 12 shows SDEnoJump can model nonlinear dynamics, while Brownian motion is far from satisfying compared to Levy processes when modeling biological systems as discussed in the introduction part. The periodic evolution procedure *G*1 → *S* → *G*2*M* → *G*1 is successfully recovered in Figure 13 by our jump diffusion model.

**Figure 11:**
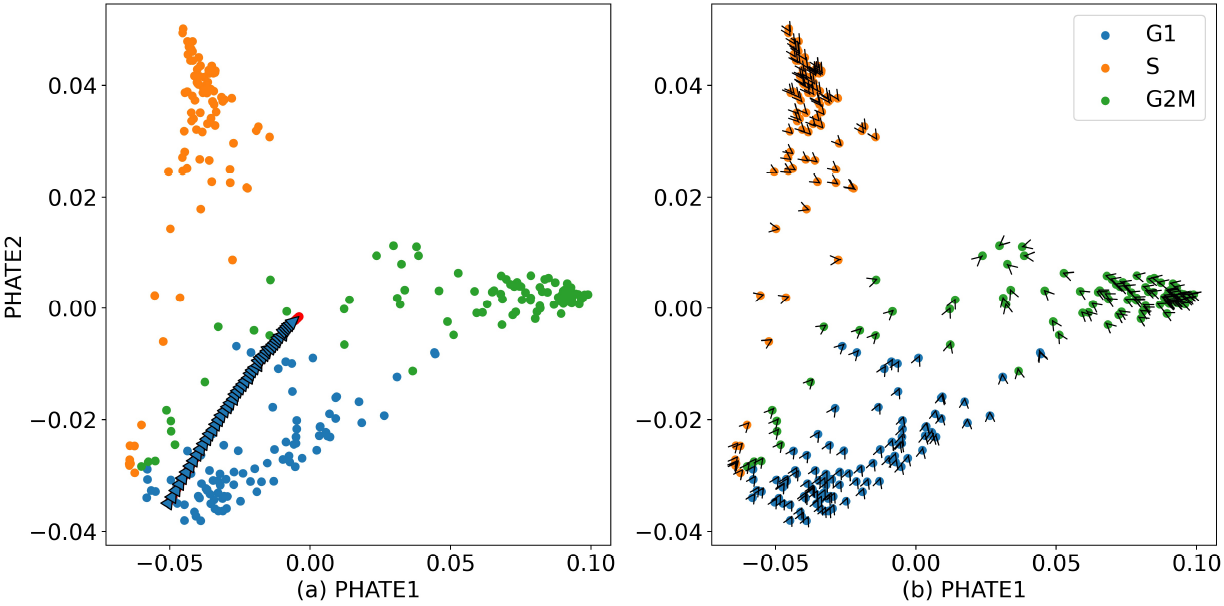
The OU process is used to model the cell cycle dataset. (a) The trajectory of a cell in two periods. (b) RNA velocity computed by the OU process for each cell.

**Figure 12:**
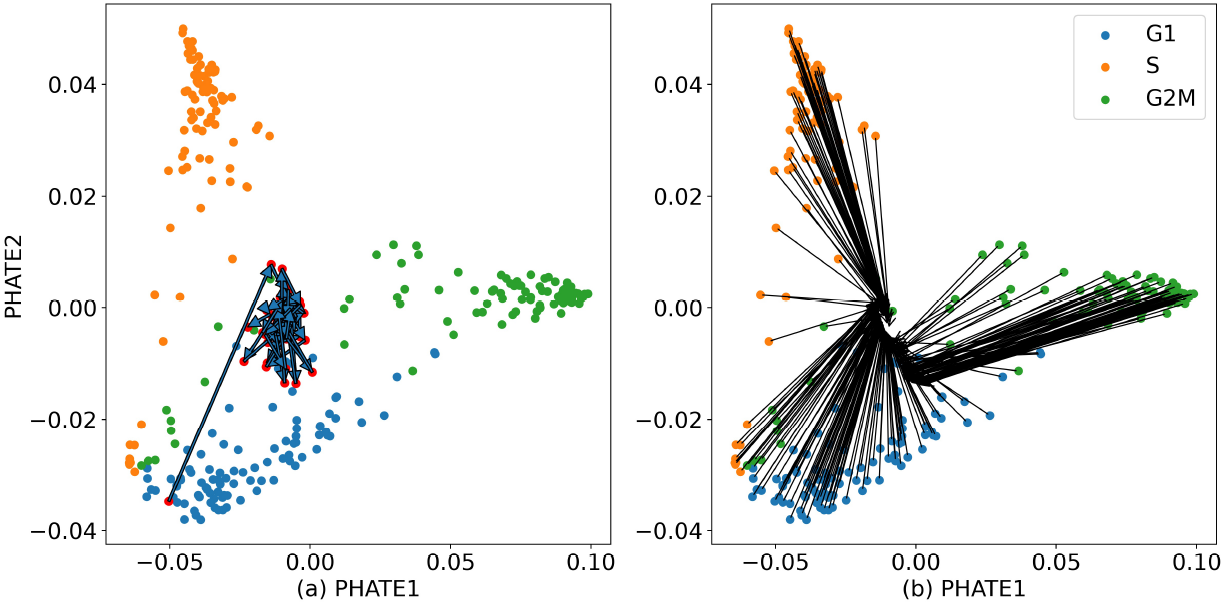
The diffusion process driven by Brownian motion is used to model the cell cycle dataset. (a) The trajectory of a cell in two periods. (b) RNA velocity computed by the diffusion process for each cell.

**Figure 13:**
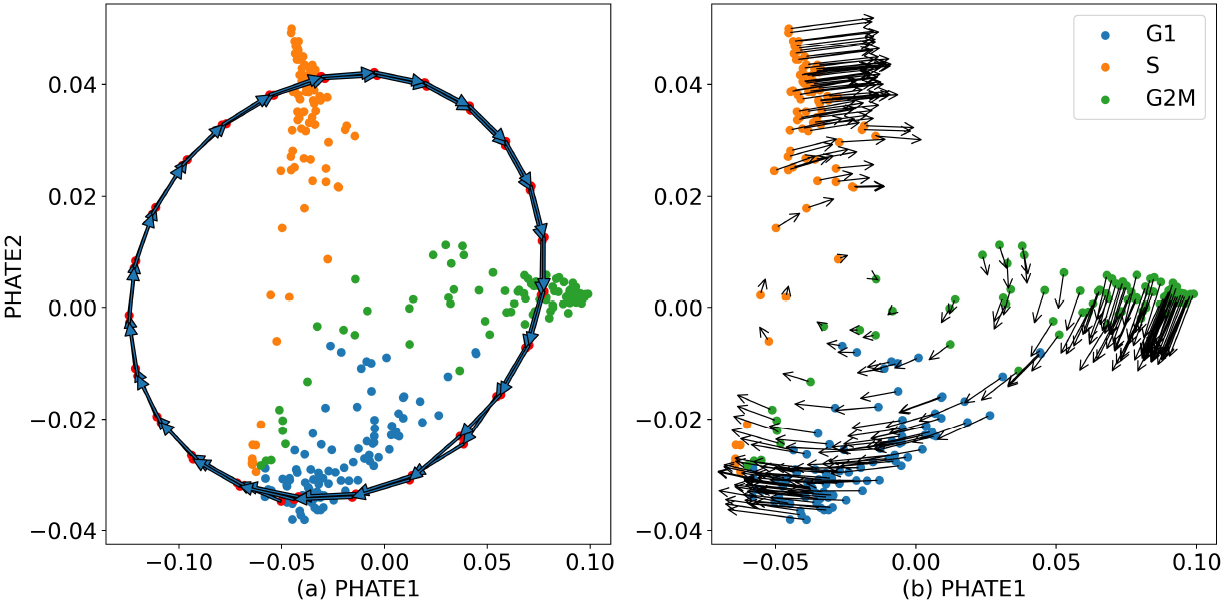
The jump diffusion process is used to model the cell cycle dataset. (a) The trajectory of a cell in two periods. (b) RNA velocity computed by the jump diffusion process for each cell.

### 3.4. Robustness analysis

We extensively test the robustness of our algorithm on hyperparameter selection including the choices of jump intensity λ, step size *δ*, evolution steps between two observation time points, and some neural network parameters. We also test the performance of our algorithm when data are perturbed. The robustness of our algorithm is confirmed in both parameter choices and perturbations of data as shown in Supplementary Note 1.

## 4. Discussion

In this paper, we propose using jump diffusion processes to reconstruct nonlinear dynamics from aggregate biological gene expression data. State variables satisfy an SDE driven by Brownian motion and a compound Poisson process, whose coefficients are set to be neural networks. The training framework for WGAN is used with *D* – 1 critics, and WGAN-div (Wu et al., 2018) is applied to implement the 1-Lipschitz constraint. The learned jump diffusion process can predict population distributions of gene expression at any developmental stage, achieve long-time trajectories for individual cells, and offer another approach to computing RNA velocity. Moreover, it gives a novel perspective to study gene expression data.

Jump diffusion processes have great advantages over the OU process and diffusion processes driven by Brownian motion in reconstructing nonlinear dynamics from aggregate data. Jumps may shorten the mean time of escaping from an attraction basin, and benefit the transitions among attractors (Zheng et al., 2016). Also, as a generalization of SDEs without jumps, our model can learn dynamics driven by Brownian motion. Experimental results show that our jump diffusion model achieves the lower Sinkhorn distances in predicting new data and succeeds in predicting the bimodal behavior of *Krt8* at D4. Also, our jump diffusion process succeeds in recovering the cell cycle dynamics. This shows our model is more suitable for biological systems.

Dimensions do not need to be too high for several top variable genes are sufficient to model the dynamic process that we are concerned with. In the stem cell differentiation dataset, 10 genes, i.e. *Krt8, Krt18, Tdh, Tagln, Mt1, Dppa5a, Mt2, Gsn, Lgals1, and Pou5f1*, are selected to learn the underlying jump dynamics. Another set of genes, i.e. *Nanog, Zfp42, Dppa5a, Sox2, Esrrb, Col4a1, Col4a2, Lama1, Lamb1, Sox17, Sparc, Krt8, Krt18, Dnmt3b*, are also used to learn a jump diffusion equation. The two learned processes predict the same trend for their commonly used genes *Dppa5a* (Supplementary Figure 5). Genes that are not included in the state variables are all treated to be noises. When dimensions are too high with small sample sizes, the learned dynamics could become quite limited.

Researchers can feed their gene expression data into our algorithm, and a jump diffusion process is achieved that best matches the observed empirical distributions. Each trajectory of the jump diffusion process represents the time evolution of a cell, and the stochasticity of trajectories determines the heterogeneity of cells. The development process of a cell is a time series, which can be clustered to describe branching dynamics. The average instantaneous rate of state change reveals the next step evolution, thus can be taken as “RNA velocity”. Moreover, in the theory of stochastic processes, a minimization of the Onsager-Machlup function using the Euler-Lagrange equation will yield the most probable path (Machlup and Onsager, 1953; Durr and Bach, 1978; Takahashi and Watanabe, 1981; Fujita and Kotani, 1982). In the future, computing quantities like attractors, critical points, and most probable paths (Chao and Duan, 2019; Wang et al., 2020; Li et al., 2021) will be the priority of next challenges.

## Acknowledgments

Our heartfelt thanks go to Prof. Minghua Deng at Peking University for offering GPU resources.

## Appendix

Proof of Theorem 1. The MSE for the estimator 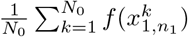 is computed by

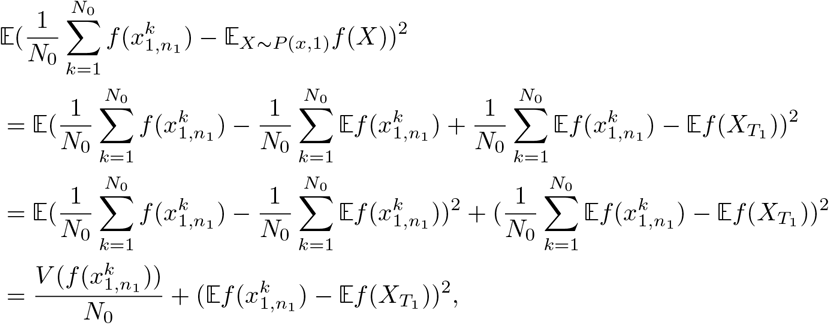

where *V*(·) denotes variance. For the second term, we have

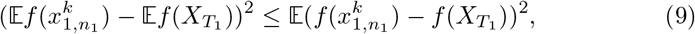

according to Jensen’s inequality. *f* is Lipschitz continuous with constant 1, so we have

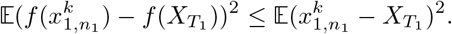

The Euler-Maruyama scheme for (3) has a strong order of convergence 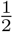, see Bruti-Liberati and Platen (2007). Therefore, we get the desired convergence rate.

